# The Impact of Small-group Education on Student Wellbeing in Undergraduate Biology: a Student Perspective on Self-Determination theory

**DOI:** 10.64898/2026.10.13.758809

**Authors:** C.M. Drion, C. Pop

## Abstract

Alarming reports about mental health have raised awareness of student wellbeing in STEM Education. Self-Determination Theory (SDT) implies that wellbeing can be improved by fulfilling the needs for autonomy, competence and relatedness. Educational design elements such as small-group teaching, peer feedback and offering a low-stakes practice environment can help fulfill these SDT needs. These elements were applied in a new academic skills course in our undergraduate Biology program. Here, we investigated how this affected the students’ sense of autonomy, competence and relatedness, and whether this was related to wellbeing. First, a survey among 48 students confirmed that student wellbeing is positively correlated to SDT fulfillment. Next, group interviews with students revealed that wellbeing is mostly dependent on relatedness, which is facilitated through small-group teaching in the first semester. Furthermore, students reported that (1) relatedness evolves through bonding with peers over the shared experience of struggle; (2) competence requires (peer) feedback and opportunities for growth, especially in laboratory coursework; and (3) a structured, low-stakes environment supports competence, but without optimal challenge, also undermines a sense of autonomy. This illustrates how students perceive the effects of small-group educational activities on their wellbeing through the lens of Self-Determination Theory.

## Introduction

In Science, Technology, Engineering and Mathematics (STEM) education, there is increasing attention for student wellbeing (Pester et al., 2023; Pfau et al., 2024). In the Life Sciences, a recent publication reported that many students experience stress, to the point where it affects their mental health (Navarro et al., 2025). Mental health is an aspect of student wellbeing, but wellbeing itself is not always clearly defined (Hossain et al., 2023). In 2015, the Program for International Student Assessment (PISA) provided a framework for the evaluation of student wellbeing, delineated in a working paper by the Organization for Economic Co-operation and Development (OECD) along 5 dimensions: cognitive, psychological, physical, social and material wellbeing (Borgonovi & Pál, 2016). Within this framework, mental health can be considered an aspect of psychological wellbeing, and at the same time, mental health is bidirectionally related to the social and cognitive dimensions of wellbeing. These domains (psychological, social and cognitive) are considered when assessing and investigating student wellbeing and it is known that they are associated with study success in higher education (Lipson & Eisenberg, 2018; van der Zanden et al., 2018). Importantly, it is also known that student wellbeing can be influenced by choices in educational design and the learning environment (Konstantinidis, 2024; Miller & Lang, 2016; Olsson et al., 2024). While effects depend largely on the specific contexts such as the type of education, year of studies, geographical location and student population (Davis & Hadwin, 2021; Dempsey et al., 2025; Kapil & Hadwin, 2026), several educational designs and teaching activities have been associated with improved student wellbeing. Especially small-group teaching activities, ranging from peer learning to tutorials in groups of 15 students and community-building activities (Jones, 2007), have been linked to improved student wellbeing (Hanson et al., 2016; Pointon-Haas et al., 2024; Springer et al., 1999; Stanton et al., 2016; Torda & Shulruf, 2021). Also, it has been shown that assessment can impact student wellbeing positively as well as negatively (Jones et al., 2021; McGuire, 2026). Interestingly, peer feedback, which is defined as ‘the learning element of peer assessment’ by Liu & Carless (2006), has been shown to impact student wellbeing positively (Ardill, 2025; Kerman et al., 2024; Topping, 2017). Moreover, it has been shown that especially in the first year of study, student wellbeing is heavily influenced by the learning environment (Knoesen & Naudé, 2018; van der Zanden et al., 2018).

In the first year of our undergraduate Biology program, we recently implemented a new course, where students meet in groups of ∼12 students and a tutor to practice academic skills in a low-stakes environment. This course aimed to engage students through peer learning activities while supporting community-building within the cohort. Because the course uses small-group teaching, peer feedback and peer learning, we suspected that the course would benefit student’s wellbeing. We set out to investigate this through a survey and group interviews, using the framework of Self-Determination theory (Ryan & Deci, 2000), which is explained below.

### Theoretical Framework

A framework to assess the psychological, social and cognitive domains of wellbeing is the Self-Determination Theory (SDT) proposed by Ryan and Deci (Ryan & Deci, 2000, 2020). According to SDT, wellbeing, motivation and optimal functioning are fostered when three psychological needs are met: autonomy, competence and relatedness. Although SDT is especially suited to predict motivation, it is a particularly useful framework to study wellbeing in the context of higher education, especially since autonomy, competence and relatedness can be influenced by the learning environment and educational design, thereby positively impacting student wellbeing (Niemiec & Ryan, 2009; Nishimura & Joshi, 2021; Zainuddin & Perera, 2017). In an educational context, autonomy is marked by a sense of agency and ownership over one’s own learning. Competence refers to the feeling of mastery, including for instance self-confidence about academic achievements. Relatedness concerns both a sense of connection, or belonging to a community, and the feeling of having support from others (Ryan & Deci, 2000, 2020). It has previously been shown that small-group teaching activities tap into autonomy, competence and relatedness (Hensley et al., 2021). For example, small-group teaching activities support the sense of connectedness (Brandl et al., 2017), and peer feedback positively impacts the sense of competence (Simonsmeier et al., 2020) and autonomy (Kerman et al., 2024).

Therefore, we hypothesized that our course would affect the sense of autonomy, competence and relatedness of students in the first year of the Biology program, and furthermore, that in our student population, SDT fulfillment (the extent to which the need for autonomy, competence and relatedness are met), is related to wellbeing.

We investigated this using a mixed methods design. We first conducted a survey to establish the relationship between wellbeing and SDT fulfillment. Based on the survey outcome, we conducted group interviews with students to explore how the components of SDT are addressed by the new academic skills course from a student perspective.

## Methods

### Teaching context

This study was conducted within the undergraduate (bachelor) program Biology at the University of Groningen in the Netherlands. The program is English-taught and consists of ∼200 students in the first year, of which ∼33% are international and ∼67% are Dutch. The newly implemented academic skills course was piloted in the academic year 2023-2024 and was subsequently embedded in the curriculum from 2024-2025 onwards. The course consists of 12 tutorials in groups of ∼12 students, led by a tutor, throughout the first semester. Students practice academic skills (reading an article, presenting, reporting research findings, creating a scientific poster), introductory lab skills (pipetting and chromatography) and professional/transferable skills (collaborating in teams, peer assessment, critical thinking and AI literacy). Assignments in the course include group assignments (scientific poster and presentation) and individual assignments (report writing), for which peer-feedback is consistently used. In addition, each group of students is assigned a mentor, who is a more senior student, to help the first-year students navigate the program as well as student life in general.

### Study design

A mixed methods design, combining quantitative survey data with qualitative focus group interviews was used to test our hypotheses. Quantitative survey data were used to test the relationship between self-reported wellbeing and SDT fulfillment (the extent to which the need for autonomy, competence and relatedness are met, according to students). The outcome subsequently informed the group interview design, which explored how students experienced those constructs in relation to the academic skills training in their first year program.

#### Survey

##### Participants

Students enrolled in the undergraduate (bachelor, BSc) Biology program at University of Groningen, the Netherlands, were invited to participate an online survey. The survey was shared with the students at the start of the academic year, in September 2025. 68 students participated in the survey (response rate ∼15%). First-year students were asked to participate in a tutorial of the first-year academic skills course, where time was allocated to filling out the survey. In addition, students in their second, third or fourth year were recruited via an announcement in the learning management system. We invited students from all years to assess wellbeing and SDT fulfillment in the entire cohort.

##### Data collection

The survey included validated questionnaires for assessing wellbeing and SDT. First, the WHO-5 Wellbeing Index was used to assess overall wellbeing over the past 2 weeks (Topp et al., 2015). To assess SDT fulfillment (the extent to which the need for autonomy, competence and relatedness are met), we used the brief SDT questionnaire, which indicates the perceived sense autonomy, competence and relatedness. An overview of the constructs, questionnaire sources and their corresponding items is provided in Table 1.

**Table 1.** Overview of the survey constructs, items, source, score computation.

| Construct + source | Items | Scale and points | Score calculation |
| --- | --- | --- | --- |
| **Wellbeing**<br>WHO-5 WellBeing Index (Topp et al., 2015) – 5 items. | Over the past 2 weeks...<br>- ... I have felt cheerful and in good spirits<br>- ... I have felt calm and relaxed<br>- ... I have felt active and vigorous (strong, energetic)<br>- ... I woke up feeling fresh and rested<br>- ... my daily life has been filled with things that interest me | 5-All of the time<br>4-Most of the time<br>3- More than half the time<br>2- Less than half the time<br>1- Some of the time<br>0- At no time | Points of all 5 items were summed and transformed to percentage scale (0-100%). |
| **Self-determination theory (SDT)**<br>(Martela & Ryan, 2024)– 3 items. | Please read the following items carefully with your BSc Biology education and personal goals in mind. Using the response scale below, indicate the extent to which you agree or disagree with that statement:<br>- I am able to do things that I really want and value in life<br>- I can do things well and achieve my goals<br>- I feel close and connected with other people who are important to me | 1 - Strongly disagree<br>2 - Disagree<br>3 - Somewhat disagree<br>4 - Neither disagree nor agree<br>5 - Somewhat agree<br>6 - Agree<br>7 - Strongly agree | Points of all 5 items were summed. |

##### Data analysis

Survey data were analyzed using IBM SPSS Statistics (v28.0.1). Composite scores were calculated for each construct according to the corresponding instrument source (see Table 1). For each score, the distribution of datapoints was tested for normality. Since the assumption for normality was violated, we tested whether there was a correlation between wellbeing score and self-determination theory score using Spearman’s. Statistical significance was determined using a two-tailed test and considered when p<0.05. Ranked data was plotted.

#### Group interviews

##### Participants

Two group interviews were conducted with five students each (N = 10 in total). Participants were recruited through an open call to all first-year students in the academic skills course, through student members of the program committee, through an advertisement in second-year courses and through the study association. We mainly focused on recruiting second year students, since they had completed the new course and were able to reflect on their entire first year of the study, but we invited students from all years to increase the diversity of the participants and explore student perceptions across the program. Participation was voluntary and independent of course assessment, survey participation or course results.

In the first interview, in December 2025, five second-year students participated. The second interview was conducted in February 2026 and here too five students participated, of which there were two first-year students, two third-year students, and one fourth-year student.

##### Data collection

Both interviews were semi-structured, guided by an interview protocol that provided structure to the discussion while allowing participants to elaborate on their own experiences and add ideas beyond the scope of the interview questions. The first interview explored which aspects, in the context of the first-year academic skills course, related to wellbeing according to students. Since the preliminary analysis of this interview, in line with the survey outcome, revealed that experiences shared by students aligned well with the components of self-determination theory, the second interview focused more explicitly on SDT components: questions addressed how students experienced the sense of autonomy, competence and relatedness components in the context first-year academic skills course.

Summaries of the interview questions (the interview protocols) can be found in the supplement, excluding the questions that spontaneously arose during the interviews in response to the discussion.

##### Data analysis

Group interview data were analyzed using thematic analysis, following an inductive-deductive approach. A directed content analysis was performed (Hsieh & Shannon, 2005) where the coding process was informed by the theoretical framework of self-determination theory, while remaining open to codes emerging inductively from the data. Coding was done by two researchers using ATLAS.ti Web (version v9.26.2). The interviews were transcribed verbatim, read and coded. Then, the interviews were re-read and codes were added, changed, or removed in consultation. The codes were categorized into three themes: relatedness, competence, and autonomy (see Supplementary Table 1). There were no other emerging themes.

## Results

### Survey findings

Respondents who completed all questions related to wellbeing were included in the quantitative analysis (n=48). The 48 survey respondents were aged 17-26 years old; 29 of the respondents identified as female, 17 as male, 2 as other, and 29 of them were Dutch. Other nationalities included Romanian, Spanish, German, as well as other nationalities (figure 1). Most respondents were of the cohort 2025 (students who started the program in September 2025). The median wellbeing score (WHO-5 Wellbeing Index) was 52 (range: 8-88). 19 out of the 48 respondents reported a wellbeing score of 50% or lower (figure 1). Spearman correlation revealed a positive correlation between the wellbeing score and the score indicating self-determination theory (SDT) fulfillment (SDT score); r_s_ (46)=0.63, P<0.001 (figure 2).

**Figure 1:**
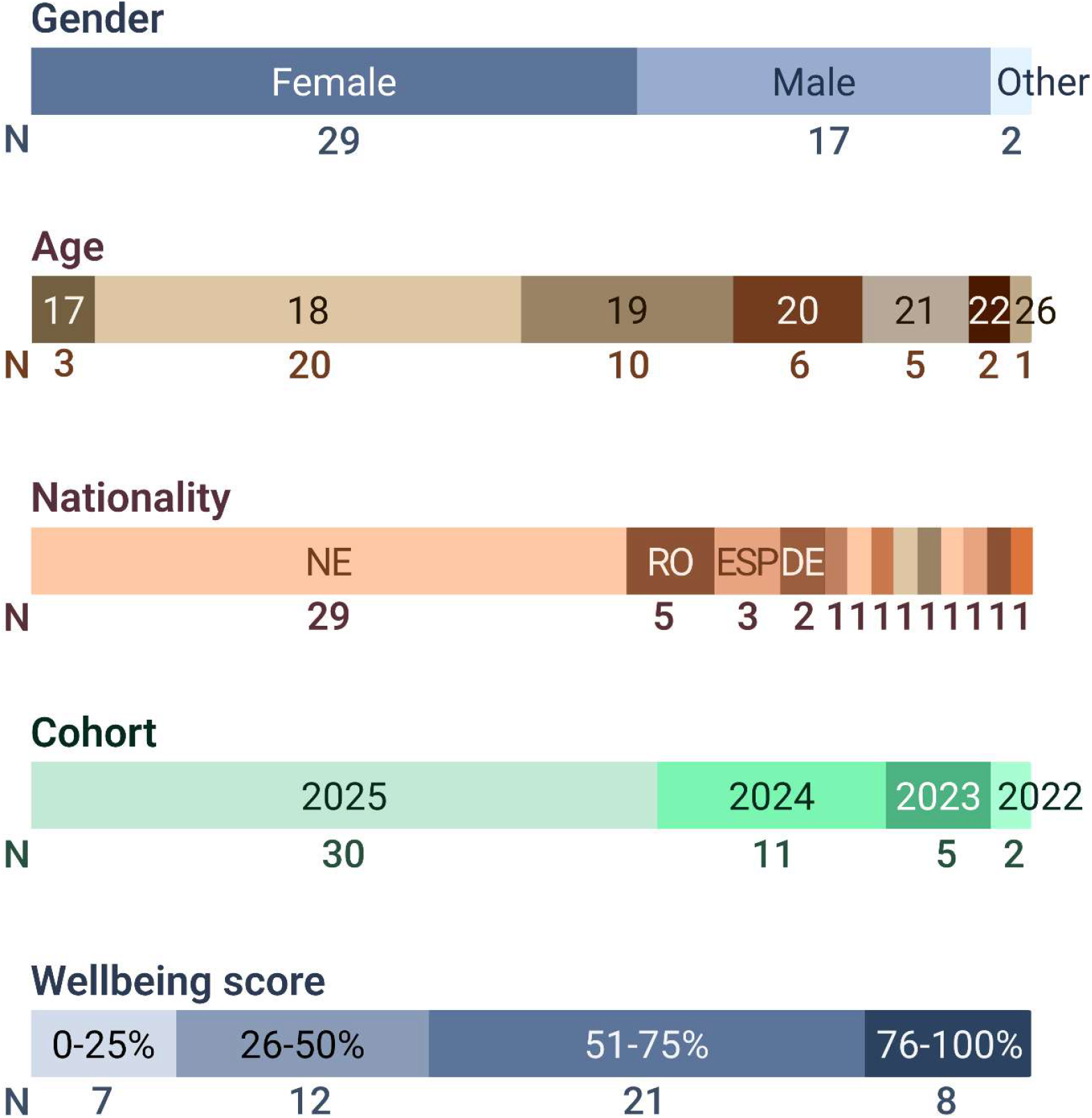
distributions of the respondents’ gender, age, nationality, cohort (starting year of the Biology program) and the wellbeing score from the WHO(5) Wellbeing Index. Nationalities: NE = Dutch, RO = Romanian, ESP = Spanish, DE = German. Nationalities with n=1: Curaçaoan /Dutch, Czech, Hungarian, Indian, Italian, Italian/American, Moldovan, Moldovan/Romanian, Ukrainian. NB. one participants did not fill in ‘Age’.

**Figure 2:**
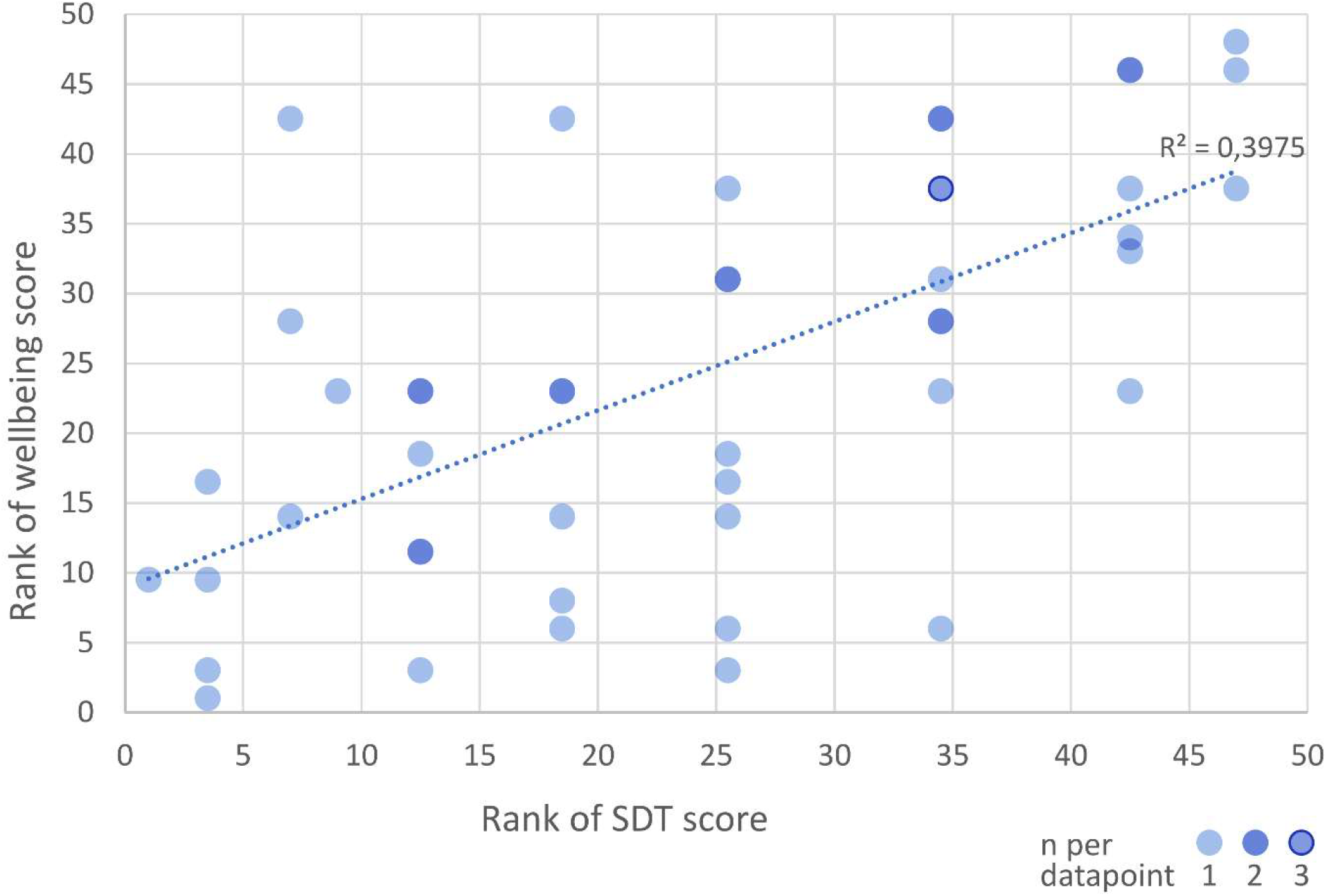
correlation between student wellbeing and self-determination theory (SDT) score (shown are ranked datapoints): r_s_ (46)=0.63, P<0.001. Darker dots indicate overlapping datapoints (see legend).

### Group interview findings

To further explore wellbeing, based on the survey outcomes, two group interviews were conducted, with 5 participants each. Through thematic analysis of the interview transcripts we coded the student quotes (see Supplementary Table 1), and subsequently grouped the codes according to themes informed by the SDT components relatedness, competence and autonomy. Most codes were grouped under the theme relatedness, followed by competence and then autonomy. Below, we explore these three themes in this order. Note that when the newly implemented academic skills course is mentioned, the course name “Basic Academic Skills” is used; abbreviated to “BAS”.

### Theme 1: relatedness

Relatedness emerged as the most prominent theme across the interviews, described by participants as foundational to their wellbeing and as a protection against stress and loneliness. Within relatedness, several subthemes emerged from the thematic analysis.

#### Small-group format as the mechanism for connection

Participants consistently linked their sense of relatedness to the small-group structure of the new academic skills course (“BAS”) and the mentor system, which reduced the anonymity of a large cohort and created repeated opportunities for contact.

> S02B [00:25:05]: “Suddenly walking into a group of 100 people, feels so overwhelming… when you’re just presented with 10 people, or in the same places you’re constantly, it’s much easier to make connections with them, make friends.”
>
> S04B [00:24:47]: “It’s nice that you see the same people over the course of the BAS course, you really feel like a class.”

These quotes illustrate the value of small-group teaching and - mentor activities to offer opportunities for students to connect with their peers.

#### Normalization of struggle through shared experiences

One of the most consistently reported effects of relatedness was the normalization of struggle through sharing experiences. Participants described how recognizing that peers faced similar difficulties helped relieve the distress associated with those difficulties, even when the underlying problem itself remained unresolved.

S05A [00:06:29]: “I think it also helps if you have a friend who’s in your study… Because you’re all going through the same thing… And you can complain about it to each other… in that complaining you find a sense of…”

S00B [00:07:03]: “if you see that others are also having those stupid little errors… then it’s like, oh, that’s just part of the process… rather than just you thinking about yourself.”

There were more quotes like these, indicating that relatedness, to these students, involved going through the same struggle and bonding over it. The bonding, but also the normalization of struggle, consequently improved their wellbeing.

#### Relatedness as a precursor to autonomy and competence

Several participants describe relatedness as foundational — a need that must be met before competence and autonomy can be pursued.

> S01B [00:02:41]: “Maybe, the relatedness is the most important in the first year… if you lack a sense of belonging at least in your first year is probably a big deal.”
>
> S02B [00:33:44]: “The lack of confidence in the social aspect kind of causes you to lack confidence in the academic part as well.”

This supports the notion that the three components of SDT are not experienced independently, but instead are interdependent.

### Theme 2: competence

Competence, the subjective experience of mastery, is best achieved in environments offering structure, feedback, the optimal level of challenge and opportunities for growth (Ryan & Deci, 2020). Some students suggested that the design of the Basic Academic Skills (BAS) course increased confidence about academic skills.

> S04B: [00:19:00] “I think skills such as reading a paper have positively impacted competence. I mean, some people do know how to do it from high school, but some people also do not know how to do it from high school, and then I think it might give you a bit more confidence, and a bit more guidance, into how to perform certain academic tasks.”

#### Peer feedback and feedback

One element that increased confidence was peer-feedback. In the next quotes, S01A and S02B refer to *providing* feedback to peers, which helped to reduce uncertainty about one’s own abilities.

> S01A: [00:33:03] “I think you could be more certain about your own abilities after the peer check than if you just hand it in and you don’t know what’s expected.”
>
> S02B: [00:19:28] “Yeah, not only reading, but writing as well. We have the diving reflex thing, and then we have the peer review thing as well, and, I mean, you saw your errors, and how people read them, but, yeah, it also gave you confidence to see, oh, you did, oh, I did well, and I can improve this, and that sort of stuff.”

Receiving feedback was indicated as essential for a sense of competence. The following account illustrates what happens when feedback is missing. After submitting an assignment without receiving any response, S06A was left uncertain whether their answers were correct:

> S06A: [00:54:37] “It’s not because I found the calculations to be unuseful or unnecessary. But combined with the fact that we have never gotten any follow-up or any feedback or any further explanation on it… when I gave it in, I wasn’t 100% sure if my answers were right. I was kind of expecting, not a grade because it was not part of the final grade, but a feedback or something. And then never got anything back from it.”

Without feedback, the task fails to help the student develop a sense of progressing competence.

#### Opportunities for growth

Feedback can promote a sense of competence, especially if growth is visible between two attempts at the same skill or learning objective. The following account about the experience of a failed exam and successful re-sit offers a concrete example:

> S04A: [00:42:12] “I think also having a re-sit helped me, because then I could kind of trace back what I did and how that apparently didn’t work. And then for the re-sit, I tried something else, and it worked, and then I kind of knew that I had to keep that up. Yep.”

The failed exam prompted the student to trace back their approach, identify what had not worked, and test an alternative. In this way, the re-sit provided the confirmatory feedback that their revised strategy worked. ‘I kind of knew that I had to keep that up,’ suggests the feedback helped enhance confidence and increased intrinsic motivation.

As part of the BAS course, students completed an introductory laboratory (‘lab’), which is the first lab they encounter in their program. The goal is to prepare them for subsequent labs, make those less daunting and offer an opportunity to practice learned skills a second time.

> S06A: [00:22:13] “…one of the reasons why I found the first lab extremely stressful was because I was learning new skills and I kind of expected myself to do it perfectly the first time. [00:23:14] And then, when it didn’t turn out the way I wanted it to, I had a very difficult time adjusting.”
>
> S06A: [00:23:28] “… compared with the second lab of the first year … I was just more accepting of changes and accepting that things will not always go the way I planned them. And then, I feel like I had a much easier time, the second time.”

S06 recounts their experience in the first laboratory course and the changes experienced over time. The initial expectations of immediate mastery of new skills meant that normal difficulty registered as personal failure and turned the learning experience into a source of distress. By the second laboratory, the student describes a shift towards acceptance of uncertainty and imperfection. Together, these accounts illustrate how opportunities for growth can promote competence development.

#### Structured environment

The Basic Academic Skills (BAS) course provides a structured environment which is, according to Ryan & Deci (2020), a beneficial factor in developing competence. Some accounts suggest that BAS provided a psychologically safer context in which students could develop a more adaptive relationship with difficulty and imperfection. For example, S06A describes how the course addressed this by providing a low-stakes learning environment in which skill practice was decoupled from summative assessment, removing the pressure:

> S06A: [00:33:31] “something that I appreciated about the tutor session was that some of the things that we did were related to also what we had in courses later. But then we first got the chance to try them out in a lower pressure environment.”

Other participants recounted the same experience:

> S01A: [00:33:51] “Yeah, I also think everything is rated very low at BAS. Like, every part is a very small part towards your final grade, which is nice, because it allows you to //S05: Make mistakes?// Yeah.”

#### Optimal challenge

The downside of a structured, low-stakes environment, however, is that it can be less challenging. A finding that emerged in the interviews concerns the degree to which activities felt relevant and appropriately challenging. While the low-stakes design of the academic skills course (BAS) was valued, participants made a distinction between activities that felt meaningfully connected to their development and those that felt restrictive or insufficiently demanding. Participants with stronger prior academic training found that the BAS activities did not build competence, but instead felt repetitive. Components of the BAS course that presented insufficient challenge to the students were completed with reluctance and lack of motivation, and were attributed little educational value.

> S03B: [00:11:00] “I think this was just, like, because I was, uh, taught in high school, so it was just a bit, like, the group was held by hand, and then you already knew what the session would be about, so it was for me then sometimes a bit useless, or not really motivating to experience plus, or learn something from it.”

S03B’s account connects this finding directly to BAS: activities already mastered in secondary education were experienced as demotivating rather than confidence-building, because they offered no meaningful challenge. There were also accounts, however, of the same activities (such as giving a presentation) being perceived as highly challenging by other students, who had had no prior training. Ideally, according to SDT, activities are designed to be optimally challenging, to maximally enable competence development. In addition, this creates an autonomy-supportive learning context, linking to the third SDT theme: autonomy.

### Theme 3: autonomy

Within the autonomy theme, several subthemes emerged, showing clear connections to competence and/or relatedness.

#### Fixed methods constrain autonomous learning strategies

A highly structured environment (described in the competence section above), can imply a trade-off with autonomy. Participants experienced constraint in their autonomy when presented with a fixed strategy to apply in an assignment. The next accounts concern the assignment where students have to develop a study strategy and discuss this with their mentors as part of the BAS course.

> S05A: [00:47:56] “I had a mentor who was like, okay, what are you going to do? And I was like, I don’t know. And she was just someone who had their life planned and everything had to go this way for you.”
>
> S01B [00:14:09]: “I also remember just, like, ‘oh, this is kind of, like, how it should be done’… I don’t want to plan things every hour, because then I feel, like, if I don’t do the thing that I was supposed to in that hour, then the next hour, I don’t feel like doing it anymore.”

S05A recounts the interaction with their mentor, who had a highly structured approach that prevented the student from exploring own ways of working. S01B experienced a similar constraint due to the fixed method approach, rather than finding it supportive. This shows that too much guidance can impair the sense of autonomy.

#### Group work reduces individual agency

In the BAS course, several assignments are group assignments, where students collaborate with one or more peer(s). Some participants identified group work as an impediment to individual agency.

> S01B: [00:11:36] “I think at least in my year it was also that there a lot of projects or group work that you had to do and that makes sense because you are bonding with the people in your mentor group. But then I felt like everything had to come back where I’m a voice of the four people, and none of the decisions that we made on those projects or assignments were mine, there was no solo assignments or something to do on your own.”

In group assignments, S01B experienced that their decisions, contributions, and learning outcomes belonged to the group rather than to themself. This illustrates the tension between relatedness and autonomy, where the former gets supported by the small-group format, and the latter undermined.

#### Mentor accessibility as autonomy support beyond BAS

The mentor relationship was valued not only for relatedness but for the sense of agency it gave students — knowing who to contact, how to access support, and feeling entitled to reach out. When this support disappeared after BAS ended, participants described a reduction in their sense of agency.

> S03B [00:35:22]: “the helping of a mentor, or a personal contact person, who was more easy to reach out, was really helpful.”

This suggests that access to an approachable, responsive support person functions as an autonomy-supportive structure, that enables students to identify problems, seek guidance, and make informed decisions.

## Discussion

This study explored student wellbeing in undergraduate Biology students through the lens of self-determination theory in the context of a newly implemented first-year academic skills course.

First, our survey data indicate that nearly 40% of the survey participants reported a wellbeing score below 50% which, according to the WHO-5 Wellbeing index (Topp et al., 2015) is an indication for testing for depression. This is in line with the national reports of the Trimbos institute in the Netherlands, which has shown that 44% of students in higher education in the Netherlands experience mental health issues like anxiety- and depressive-like feelings (Nuijen et al., 2025). Although this is a slight decrease compared to the report of 2023, which was right after the COVID pandemic (Nuijen et al., 2023), it is still a high number, which warrants further research into student wellbeing.

Our survey findings indicate that student wellbeing was correlated to fulfillment of Self-Determination Theory (SDT) needs: autonomy, competence and relatedness (Ryan & Deci, 2000, 2020). Based on these outcomes, we conducted group interviews with students to further explore these needs and investigate how, according to the students, the new academic skills course influenced those needs and in turn affected their wellbeing. We were able to identify both positive and negative effects of the course design on autonomy, competence and relatedness, and how these needs are perceived by students.

### Relatedness

The relatedness need of Self-Determination Theory was indicated by the students as most important for their wellbeing. Relatedness is sometimes referred to as a ‘sense of belonging’, of which the importance for student wellbeing has been highlighted by previous research (Romeo et al., 2024). The 2025 Trimbos report (Nuijen et al., 2025) also indicates that students experiencing a sense of belonging had better mental health. Interestingly, interview outcomes suggested that the sense of belonging was not identified by feeling a connection to the field of study. Instead, students associate this with connecting to other students over the shared experience of struggle. ‘Struggle’ in our interviews referred to failed exams, busy schedules and performance anxiety. Sharing experiences about this led to enhanced feelings of relatedness, which the students indeed indicate as critical for their wellbeing.

### Competence

Additionally, this shared experience also contributed to the normalization of struggle. In line with this, a previous study found the normalization of struggle an important factor for wellbeing in STEM PhD students (Posselt, 2018). It also coincides with the findings of Volstad et al. (2020), who found that considering ‘challenge’ a normal part of life when transitioning from high school to university is an indicator for ‘flourishing’ in first-year students. In Biology students it has been shown that the ability to overcome struggles influences performance (Tracy et al., 2022). This connects to the experience of one of our interviewed students, who experienced their first lab as ‘very stressful’ but by working through this struggle, had a much better experience in the second lab. This shows how offering two instances of educational activities with the same learning objective creates an opportunity for growth. This is in line with Ryan and Deci, who mention this as an important factor for competence (Ryan & Deci, 2020). From our interviews it also became apparent that students were more accepting of a struggle in a low-stakes environment, like the Basic Academic Skills (BAS) course. The course was intentionally designed to allow for trial and error, in a small group of peers with the same tutor: a familiar and comfortable setting. Assignments were generally not perceived as challenging. Although this helped to build confidence in some students, it did undermine motivation in others, which shows that striking the right balance between a safe environment and an optimal challenge is crucial. A strategy to deal with this is to offer students a choice in the type or level of an assignment to complete, which enhances both their sense of competence and autonomy (Evans & Boucher, 2015; Hensley et al., 2021). Further, students confirmed that peer feedback activities did increase self-confidence, and thereby, their sense of competence, which is in line with previous reports (Kerman et al., 2024; Simonsmeier et al., 2020).

### Autonomy

Autonomy requires an environment that offers (some degree of) freedom, and the structured environment of the BAS course in some instances led to a diminished sense of autonomy. Yet, autonomy also requires a sense of agency. Interestingly, the interview data suggest that agency was increased by access to support such as a mentor which is consistent with previous studies (Hamilton et al., 2019; Kayvani et al., 2026; Lindsay et al., 2023).

### Limitations

First of all, this study was conducted in a confined context of first-year Biology students at a Dutch university. Although the group of participating students of students represents our general student population well, we acknowledge that our sample is restricted to students who were willing to participate. Second, we used brief surveys: for the Self-Determination Theory (SDT) fulfillment, the score consisted of points from three questions only: one per domain (autonomy, relatedness, competence). Although more extensive surveys are available, we chose the short version to maximize the response rate. Third, we held two separate group interviews with slightly different approaches. However, thematic analysis was applied similarly for both transcripts, and the combined outcomes could therefore be used to compile a joint dataset. Moreover, direct quotes were used to maximize transparency about the outcomes. Fourth, we did not correct for age, gender, nationality or cohort when analyzing the correlation between wellbeing and SDT fulfillment – this was intentional, because the study aim was to investigate wellbeing in a representative sample of our Biology student population. Comparison of wellbeing scores between Dutch and international students, and between male and female respondents, did not produce significant differences.

Lastly, The SDT framework is not supported unanimously as a tool for assessing wellbeing. It has been critiqued (Trigueros et al., 2019) and also, SDT elements have been shown to be highly context-dependent (Dempsey et al., 2025; Nishimura & Joshi, 2021). Still, although SDT mainly addresses intrinsic motivation (Grenier et al., 2024) it is nevertheless a strong indicator for wellbeing.

## Conclusions

In conclusion, our findings provide a student perspective on the effects of small-group teaching on student wellbeing through the lens of Self-Determination Theory (SDT). Integrating the findings of the survey and the group interviews, we conclude that relatedness, autonomy and competence are evident elements of wellbeing for first-year Biology students. Further, these elements can be impacted by educational design, especially through small-group teaching, (peer) feedback activities and a low-stakes learning environment. Notably, implementing educational innovations requires balancing the different SDT needs, for instance when providing structure and a low-stakes environment, which promotes competence but undermines autonomy. It is therefore essential to actively and thoroughly explore the student perspective on how educational design affects their wellbeing. Taken together, our conclusions support the need for small-group education to promote student wellbeing STEM education.

## Acknowledgements

This study was funded by the Scholarship of Teaching and Learning (SoTL) grant (2025) of the Teaching Academy Groningen.

We thank Dr. Salome Scholtens for advising us on the surveys and sharing her insights, and Dr. Sofie van den Eynde, Dr. Miguel Pérez-Torres and Dr. Edith Koh for advising us on the data analysis and writing.

## Ethics statement

This research was conducted in adherence to the Ethical Guidelines for Educational Research, Fifth Edition (BERA, 2024). The study was reviewed and approved by the Research Ethics Committee of the Faculty of Science and Engineering, University of Groningen (Reference ID: FSE.EC25014, approval date July 17, 2025). All participants provided informed consent (written for interviewees and via an online consent form for survey respondents) prior to their participation. Participants were informed of the study’s purpose, their right to withdraw at any time without penalty, and the measures taken to ensure confidentiality and anonymity of their responses.

