## Supplement for "The Impact of Small-group Education on Student Wellbeing in Undergraduate Biology: a Student Perspective on Self-Determination theory"

Contents


- Interview protocols
  - December 2025
  - February 2026
- Supplementary table 1

Interview protocols

Interview protocol - December 2025

1. What do you consider wellbeing in the context of your studies?
2. In the context of the previous question (wellbeing in study context): how do you look back to your first semester (so far)?
    a. Which factors have positively affected wellbeing?
    b. Which factors have negatively affected wellbeing?
    c. (How) did your wellbeing change over the course of your studies?

3. How do the following aspects influence wellbeing, in your view?

    a. Self-management (how well are you managing your own future, priorities, social contacts, decisions, life in general)
    b. Self-reflection (considering your own beliefs, values, acting, thinking)
    c. Self-determination (being able to do the things you value in life, achieve your goals, feeling close to people important to you)
    d. Sense of belonging (connected to community, supported by mentors/friends)
    e. Self-confidence about what is expected and your own competence

4. In relation to Basic Academic Skills (all have experience with the course?)
    a. Which factors related to BAS have positively impacted wellbeing
    b. Which factors related to BAS have negatively impacted wellbeing

5. How have the following aspects changed over the course of your studies?

    a. Self-management (how well are you managing your own future, priorities, social contacts, decisions, life in general)
    b. Self-reflection (considering your own beliefs, values, acting, thinking)
    c. Self-determination (being able to do the things you value in life, achieve your goals, feeling close to people important to you)
    d. Sense of belonging (connected to community, supported by mentors/friends)
    e. Self-confidence about what is expected and your own competence

Interview protocol – February 2026

Introduction to Self-determination Theory
autonomy - the need to have freedom over your choices, to make decisions of your own volition
competence - the desire to feel effective, to feel like you are capable of performing tasks, adapting, learning, as opposed to feeling helpless
relatedness - the desire to feel connected with others and to have meaningful relationships

Goal = to find out IF and HOW BAS influences these components (and with that, wellbeing)

1. How do the following aspects influence wellbeing, in your view?
    - Autonomy
        - Self-management (how well are you managing your own future, priorities, social contacts, decisions, life in general)
        - Self-reflection (considering your own beliefs, values, acting, thinking)

    - Relatedness
        - Sense of belonging (connected to community, supported by mentors/friends)

    - Competence
        - Self-confidence about what is expected and your own competence

2. In relation to Basic Academic Skills:
    - Which factors related to BAS have positively impacted autonomy?
    - Which factors related to BAS have negatively impacted autonomy?
    - Which factors related to BAS have positively impacted competence?
    - Which factors related to BAS have negatively impacted competence?
    - Which factors related to BAS have positively impacted relatedness?
    - Which factors related to BAS have negatively impacted relatedness?

3. Wellbeing: what is wellbeing to you and how has this changed over the first semester / did BAS affect this?

Supplementary table 1: codes organized per theme

| Theme | Code name | Code definition |
|---|---|---|
| autonomy | structure/discipline | externally imposed structure supports discipline |
| | personal relevance | the degree to which students perceive a task as meaningful and worthwhile in relation to their own goals, interests, or personal growth |
| | deadlines & evaluations | the presence of time constraints and formal assessments that structure academic performance, which can act as sources of pressure, motivation, or anxiety |
| | sense of individuality | the awareness and appreciation of one's unique qualities, experiences, and perspectives, distinct from those of peers or the group (Bornholt, 2000) |
| | optimal challenge | when tasks are at or slightly above a student's ability level, it facilitates intrinsic motivation |
| | agency | sense of being in control of your own actions (self-efficacy, self-steering) |
| | extrinsic pressure | the presence of factors that influence motivation and performance, that come from outside the individual |
| competence | study-life balance | the ability to maintain a healthy equilibrium between academic responsibilities and personal life |
| | stress factors | identified stressors |
| | accomodation period | students mention having difficulties at the beginning of their studies that eventually went away as they were accomodating |
| | feeling of falling behind | the perception that one is not keeping up with peers, expectations, or the pace of the course, often accompanied by anxiety or self-doubt |
| | managing expectations/intrinsic pressure | the process of aligning one's anticipations about performance, outcomes, or experience, to align with reality |
| | self-regulation (resilience) | the ability to manage one's own thoughts, emotions and behaviours in response to challenging situations |
| | feeling of achievement | a sense of accomplishment that arises when one is completing a task |
| | feedback | receiving information about one's performance |
| | clear expectations | knowing in advance what is required or anticipated in a given task or situation |
| | confidence | a general sense of trust in one's own abilities and judgment, connected to self-esteem |

| | | |
|---|---|---|
| | safe learning environment | a low pressure space where students can practice without consequences |
| | self-efficacy | the feeling of being capable of managing tasks, challenges, or situations independently, without relying on external support |
| | adaptability | the ability to adjust strategies in response to changing situations |
| | competence | the feeling of being capable and effective in carrying out tasks, related to performance and ability in a certain domain |
| | self-esteem | confidence in one's own worth, abilities, or morals; self-esteem encompasses beliefs about oneself as well as emotional states, such as triumph, despair, pride, and shame (Hewitt, 2009) |
| relatedness | support system | the network of people and resources that provide assistance academically or emotionally |
| | sense of belonging | the feeling of being accepted, included, and valued as part of a group (Pedler et al, 2021) |
| | social comparison | assessing ones abilities and worth against others |
| | shared experience | the knowledge that others have similar experiences brings comfort |
| | mediate connection | processes that facilitate students in forming bonds with peers |
